## Supplementary Information 1 for "Genome assembly of the milky mangrove *Excoecaria agallocha*": EBPHK_Mangrove Plant Supplementary Information.v1.docx

**Supplementary Information 1.** Summary of genomic sequencing data.

| **Liabrary** | **Reads** | **Bases** | **Coverage(X)** | **Accsion number** |
| --- | --- | --- | --- | --- |
| PacBio HiFi | 3,503,202 | 33,204,508,502 | 25 | SRR24631716 |
| Omnic | 500,462,964 | 75,069,444,600 | 56 | SRR26908863 |

**Supplementary Information 2.** Genome assembly QC and contaminant/cobiont detection.

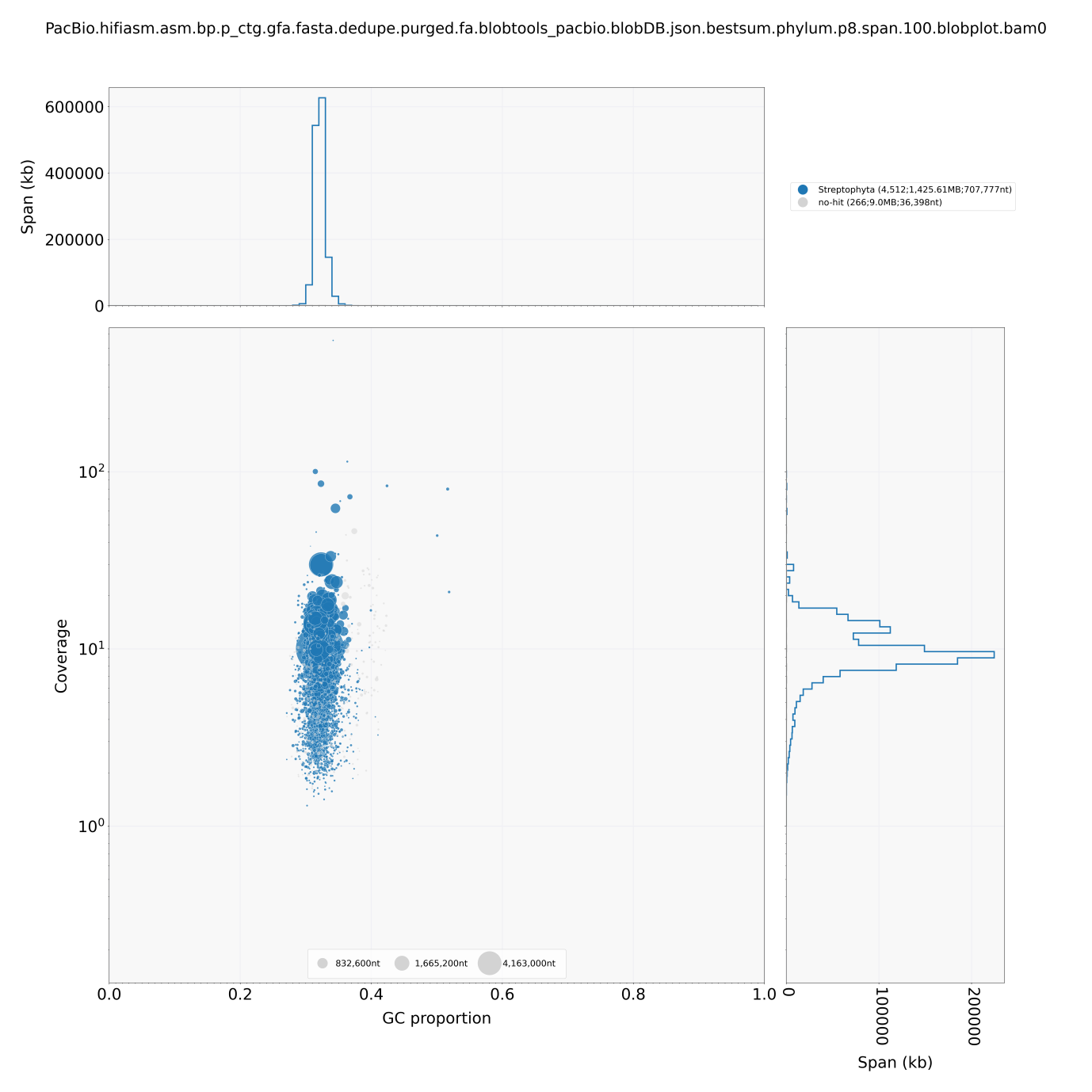

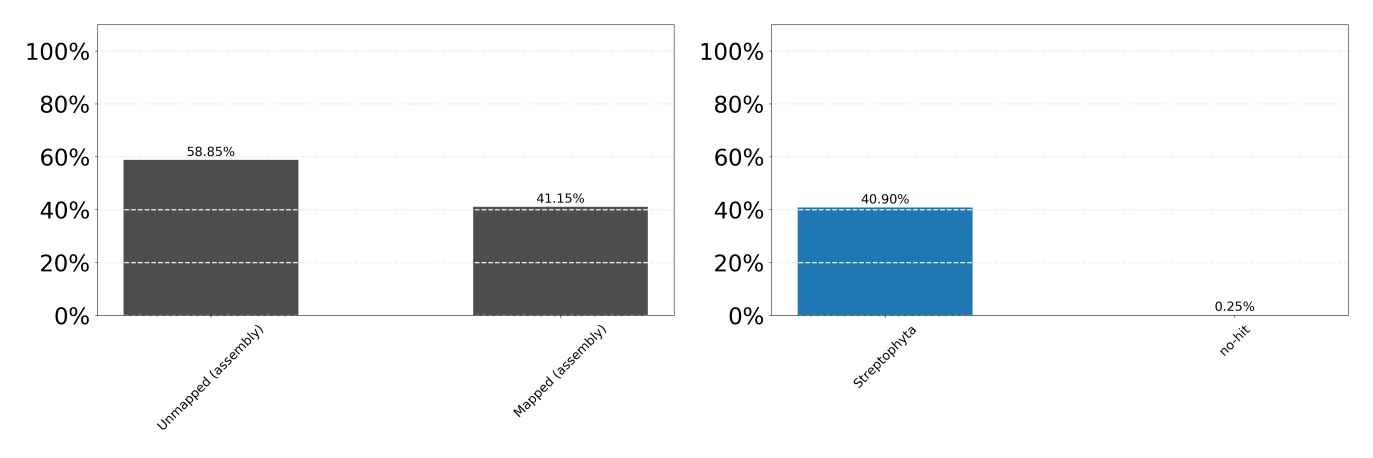

**Supplementary Information 3.** Information of 18 pseudochromosomes and BUSCO result.

| Chr Number | scaffold_length | scaffold_id | % of whole genome |
| --- | --- | --- | --- |
| 1 | 126,146,454 | scaffold_2_1 | 9.47% |
| 2 | 125,683,813 | scaffold_3_1 | 9.43% |
| 3 | 115,235,106 | scaffold_1_1 | 8.65% |
| 4 | 75,925,046 | scaffold_4_1 | 5.70% |
| 5 | 71,251,965 | scaffold_5_1 | 5.35% |
| 6 | 70,346,551 | scaffold_6_1 | 5.28% |
| 7 | 59,451,589 | scaffold_7_1 | 4.46% |
| 8 | 58,948,931 | scaffold_8_1 | 4.42% |
| 9 | 57,619,229 | scaffold_9_1 | 4.32% |
| 10 | 52,150,882 | scaffold_10_1 | 3.91% |
| 11 | 48,057,327 | scaffold_11_1 | 3.61% |
| 12 | 46,431,560 | scaffold_12_1 | 3.48% |
| 13 | 43,065,745 | scaffold_13_1 | 3.23% |
| 14 | 41,974,720 | scaffold_14_1 | 3.15% |
| 15 | 41,514,226 | scaffold_15_1 | 3.12% |
| 16 | 40,341,847 | scaffold_16_1 | 3.03% |
| 17 | 37,296,423 | scaffold_17_1 | 2.80% |
| 18 | 35,487,587 | scaffold_18_1 | 2.66% |
| SUM: | 1,146,929,001 | | 86.08% |
| BUSCO: | C:92.7%[S:35.3%,D:57.1%] | | |
